# Social Regulation of Egg Size Plasticity in the Honey Bee is Mediated by Cytoskeleton Organizer Rho1

**DOI:** 10.1101/2022.05.22.492980

**Authors:** Bin Han, Qiaohong Wei, Esmaeil Amiri, Han Hu, Lifeng Meng, Micheline K. Strand, David R. Tarpy, Shufa Xu, Jianke Li, Olav Rueppell

## Abstract

Egg size plasticity represents an adaptive reproductive strategy in numerous organisms, including the honey bee, *Apis mellifera*. However, the proximate causation of this plasticity and egg size in general is unknown. We show that honey bee queens predictably and reversibly adjust egg size in response to their colony size and that this plasticity is an active response to the queens’ perception of colony size instead of a consequence of egg laying rate. The egg size increase involves changes of 290 ovarian proteins, mostly related to increased energy metabolism, protein transport, and cytoskeleton functions. Spatio-temporal expression analysis of the small GTPase *Rho1* indicates its central role in egg size regulation, which we confirm by RNAi-mediated gene knock-down and expression analyses. The molecular adjustments that promote maternal investment of honey bee queens in response to their social environment thus reveal a novel mechanism of egg size regulation.

## Introduction

Life history evolution involves trade-offs among numerous traits ^1–4^ but the underlying mechanisms are often unclear. Optimization of offspring provisioning has resulted in a wide variety of reproductive strategies that are characterized by species-specific trade-offs between offspring size and number. Numerous studies have analyzed the trade-off between offspring number and size across and within many different species ^5–8^, with the notable exception of social insects. Offspring provisioning results in inter-generational effects that can profoundly affect organismal phenotypes and evolutionary dynamics ^9–11^. Environmental conditions typically lead to plastic responses, often in the form of variation in offspring number. However, propagule size can also be adjusted, particularly in plants, insects ^6^ and some bird species ^12^. While the adaptive reasons for egg size variation have been studied extensively ^7,13^, little is known about the proximate regulation of egg size, which is equally important for understanding this fundamental life-history trait ^2,14^.

Social evolution changes selection pressures and adaptive evolution due to kin selection ^15^, particularly in eusocial insects with colonies that form a distinct level of selection ^16^. In these species, many individuals contribute to a homeostatically regulated colony environment ^17^, pronounced phenotypic plasticity results in individuals specialized for particular functions ^18^, and resource transfers among kin influence reproductive value ^19^. Thus, life history evolution in eusocial insects differs fundamentally from that of other species ^20,21^ and has generated some extraordinary trait combinations that defy traditional life-history trade-offs ^22^. Specifically, the reproductively specialized queen caste is typically well-provisioned and cared for by non-reproductive workers, which also perform all of the intensive brood care that is characteristic for eusocial insects. As such, social insect queens may not be resource-limited despite their very high reproductive effort ^23^. Nevertheless, honey bee (*Apis mellifera*) queens display plasticity in egg size that is consistent with patterns in solitary species and corresponds to an adaptive investment hypothesis ^24^: Egg size is relatively small under favorable conditions, such as food abundance and a large colony (>8,000 workers), but is relatively large when food availability or colony size decline ^25^. A honey bee queen typically serves as the sole reproductive in her colony and is fed and cared for by her workers. However, it is unknown how much food she receives and how queen care changes with colony size. Queens can be experimentally transferred between colonies, although some are rejected and killed. Queen condition also affects egg size ^26^, and egg size differs between worker- and queen-destined eggs ^27^ with important consequences for caste determination ^28^. However, none of these phenomena have been explored further to understand the underlying mechanisms that governs variation in egg size.

Here, we report our findings of an in-depth investigation of how egg-size plasticity in honey bee queens is regulated and contribute knowledge of the molecular control of insect egg size in general, which has been difficult to determine because hundreds of genes may be involved ^14^. Our results demonstrate that the previously identified effects of colony size on queen egg size ^25^ are reversible and not fixed. We further establish that egg size is actively regulated and not a passive consequence of egg laying rate, with larger eggs produced in smaller ovaries. We further show that the social cue triggering changes in egg size within the queen does not require physical contact. Proteome comparisons between queen ovaries producing small versus large eggs indicate a central role of protein localization and cytoskeleton organization. We additionally demonstrate that the knock-down of the central cytoskeletal regulator *Rho1* significantly decreases egg size. Our data thus suggest that social cues are translated into specific molecular processes to control plastic reproductive provisioning, which presumably evolved as an adaptation to the colonial life cycle of honey bees but maybe more generally applicable to other oviparous animals.

## Results

### Honey bee queens reversibly adjust egg size in response to colony size

The first experiment involved repeated transfers of queens among colonies of different sizes, which was designed to expand our previous findings that honey bee queens can regulate their egg size in response to colony conditions. Sister queens that were housed in medium-sized colonies at the start of our first experiment produced a range of intermediate egg sizes with significant inter-individual differences (F_(10,219)_ = 31.5, *p* < 0.0001). Over the course of the first week, egg sizes significantly increased (paired t-test t = 5.7, df=10, *p* < 0.001). The first and second measurements were highly correlated (R_P_ = 0.80, n = 11, *p* = 0.003), indicating consistent differences among queens. After transfer from medium to small colonies, the egg size of all six queens increased significantly (for each queen: F_(1,38)_ = 23.7 to 153.3, *p* < 0.001). In contrast, egg size significantly decreased for all five queens that were transferred from medium to large colonies (F_(1,38)_ = 8.9 to 53.2, all *p* < 0.005). Our reciprocal transfers after the fourth week showed that egg size adjustments were reversible because all three queens successfully transferred from large to small colonies significantly increased their egg sizes (F_(1,38)_ = 143.8 to 1001.8, all *p* < 0.001) and all five queens transferred from small to large colonies significantly decreased the size of their eggs (F_(1,38)_ = 123.0 to 699.4, all *p* < 0.001). The egg size of most queens did not significantly change between separate measures in the same-sized colonies (3^rd^ versus 4^th^ or 5^th^ versus 6^th^ week). Thus, honey bee queens consistently adjust the size of their eggs in response to colony size despite inter-individual differences in absolute size (Fig. 1, Table S1).

**Fig. 1.**
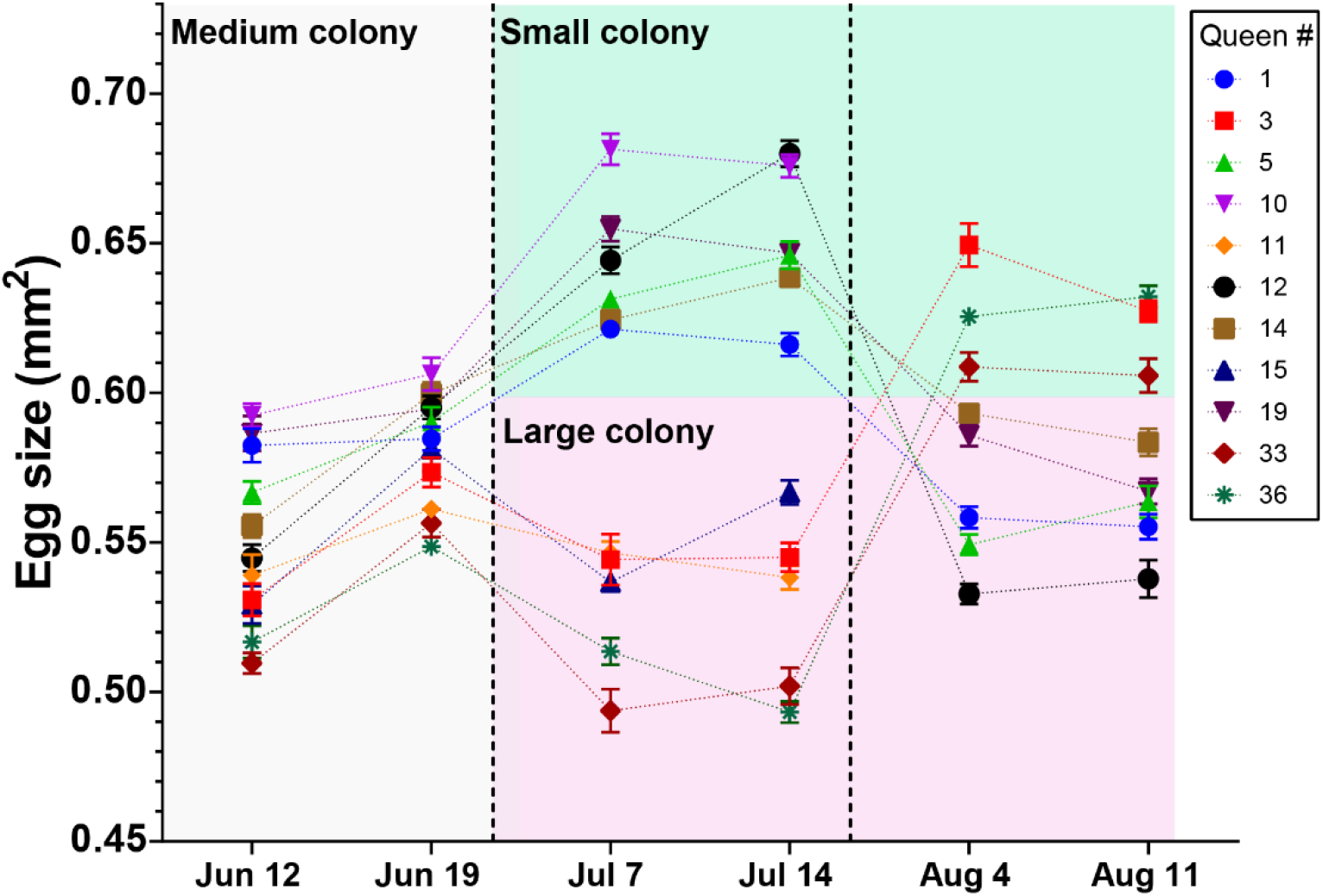
Honey bee queens reversibly adjust egg size according to colony size. The egg size (n = 20 for each data point) of individual queens (unique color symbols) was measured for six weeks while they were moved from medium to small to large or from medium to large to small colonies. Despite the presence of individual and environmental differences, these experiments show a strong and consistent negative relation between egg size and colony size.

The surviving queens of this experiment, plus two additional large-egg producing queens to increase sample size, were compared with regard to size, body weight, and ovary weight. Queens in small colonies had significantly lighter ovaries than those in large colonies (F_(1,8)_ = 10.2, *p* = 0.013), while body size (F_(1,8)_ = 0.3, *p* = 0.596) and wet weight (F_(1,8)_ = 0.8, *p* = 0.402) did not differ (Fig. 2A and Table S2). These results were confirmed in a second comparison among queens housed in small and large colonies (ovary: F_(1,6)_ = 28.7, *p* = 0.01; size: F_(1,6)_ = 0.07, *p* = 0.805; weight: F_(1,6)_ = 0.3, *p* = 0.627; Fig. 2B and Table S2), while a third data set indicated that similar-sized queens (F_(1,10)_ = 0.1, *p* = 0.748) housed in small and large colonies can differ not only in ovary weight (F_(1,10)_ = 18.5, *p* = 0.002) but also in body weight (F_(1,10)_ = 5.6, *p* = 0.039; Fig. 2C and Table S2).

**Fig. 2.**
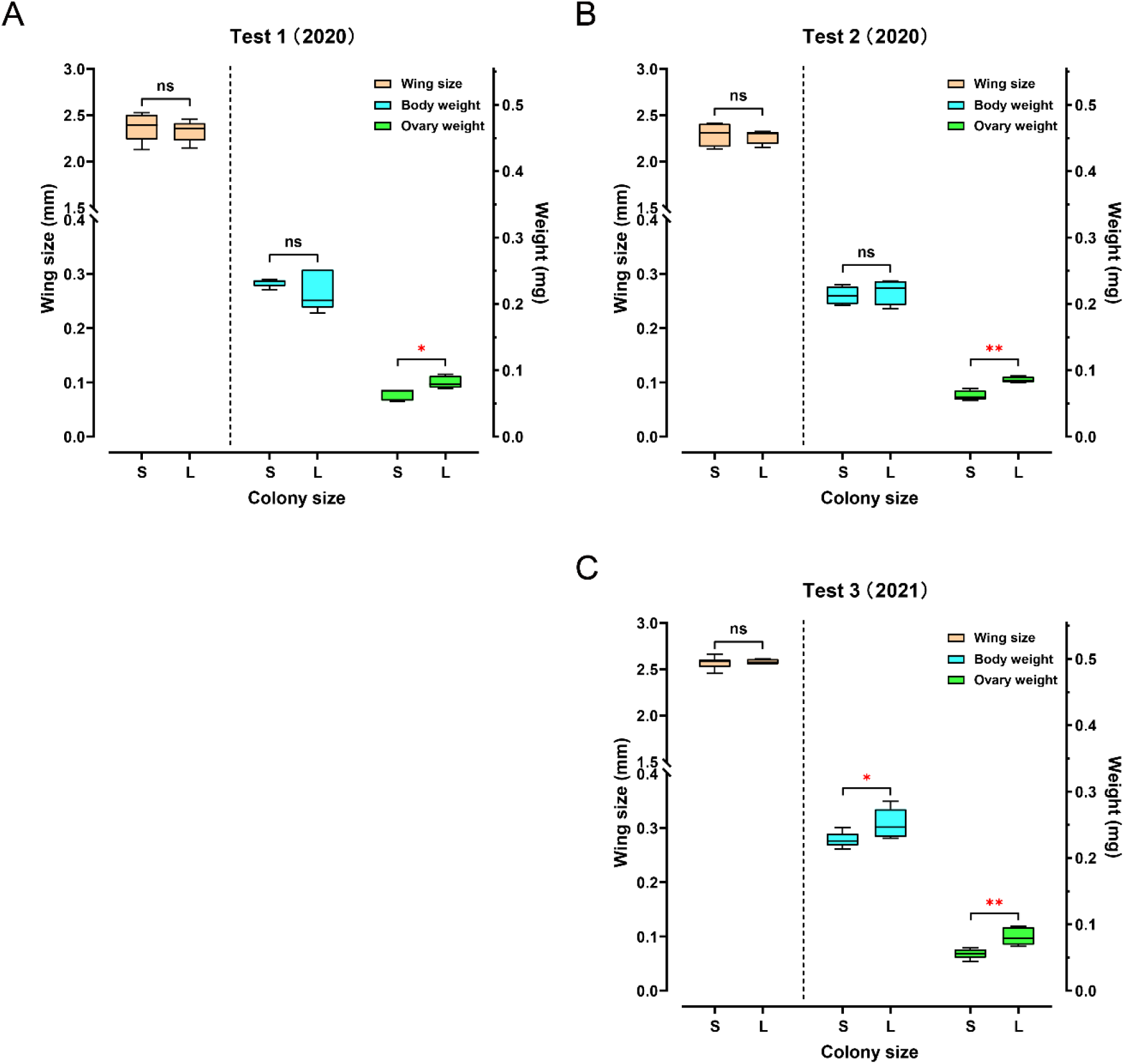
The queens’ ovary weighs less in “Small” colonies than in “Large” colonies. While queen size, measured as wing size, was not significantly different between queens in “Large” (L) and “Small” (S) colonies, ovaries were consistently lighter in queens from small colonies than in queens from large colonies. Total body weight of queens showed no significant difference between the two groups in the first **(A)** and second **(B)** experiments, but queens in large colonies were significantly heavier than queens in small colonies in the third experiment **(C)**.

### Egg size is unaffected by egg-laying rate

To test whether small egg size is merely a passive consequence of high egg-laying rate, we measured the effect of oviposition restriction on egg size: Queens from large colonies were caged to restrict oviposition and decouple egg-laying rate from colony size. None of the restricted queens significantly increased her egg size (F_(1,38)_ = 0.02 to 1.8, all *p* > 0.1). Queens in an unmanipulated control group during the same time did not change egg size either (F_(1,38)_ = 0.005 to 0.6, all *p* > 0.4), and egg sizes were similar between the restricted and unrestricted groups overall (Fig. 3 and Table S3).

**Fig. 3.**
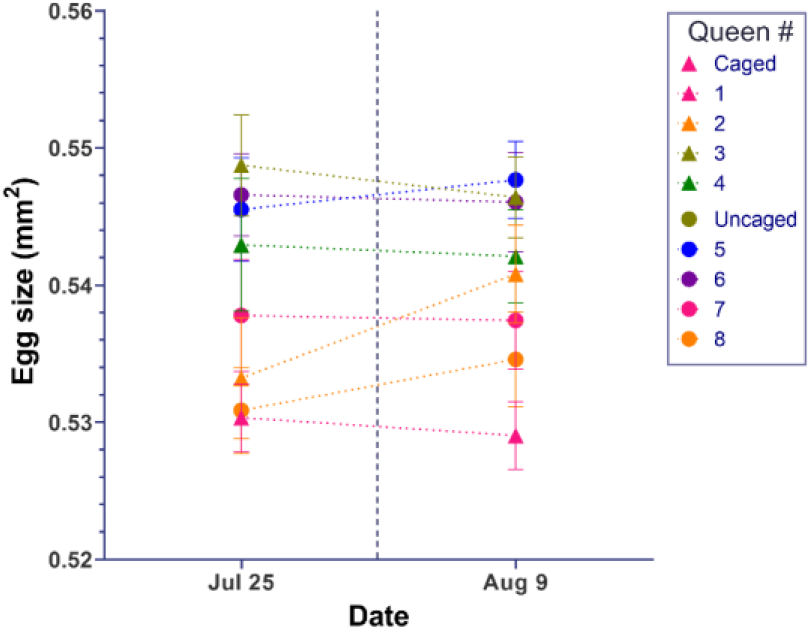
Egg size of queens is not affected by egg-laying rate. After egg size of individual queens in large colonies was measured, treatment queens (triangular symbols) were confined on capped brood comb that did not allow any oviposition while the control queens (circle symbols) had free access to empty comb for oviposition. After 14 days, the egg size in neither group of queens changed significantly. Individual means ± S.D. are shown.

### Queens adjust their egg size in response to perceived instead of actual colony size

To better understand how colony size influences queen egg size regulation, the perceived but not the physical colony size of small colonies was extended. The queens in “Small” colonies, producing relatively large eggs, were paired via a double-screened tunnel with medium-sized hive boxes that either contained empty frames or a queenless, “Medium” colony. All three queens paired with a regular colony reduced the size of their eggs compared to their initial egg size (Q1: F_(3,76)_ = 34.5, *p* < 0.001; Q2: F_(3,76)_ = 42.5, *p* < 0.001; Q3: F_(3,76)_ = 14.6, *p* < 0.001; post-hoc tests indicated significant differences only between measurements before and after manipulation; Fig. 4 and Table S4). In contrast, none of the three control queens significantly changed their egg size during the experimental period (Q1: F_(3,76)_ = 1.3, *p* = 0.297; Q2: F_(3,76)_ = 1.6, *p* = 0.196; Q3: F_(3,76)_ = 1.0, *p* = 0.379; Fig. 4 and Table S4).

**Fig. 4.**
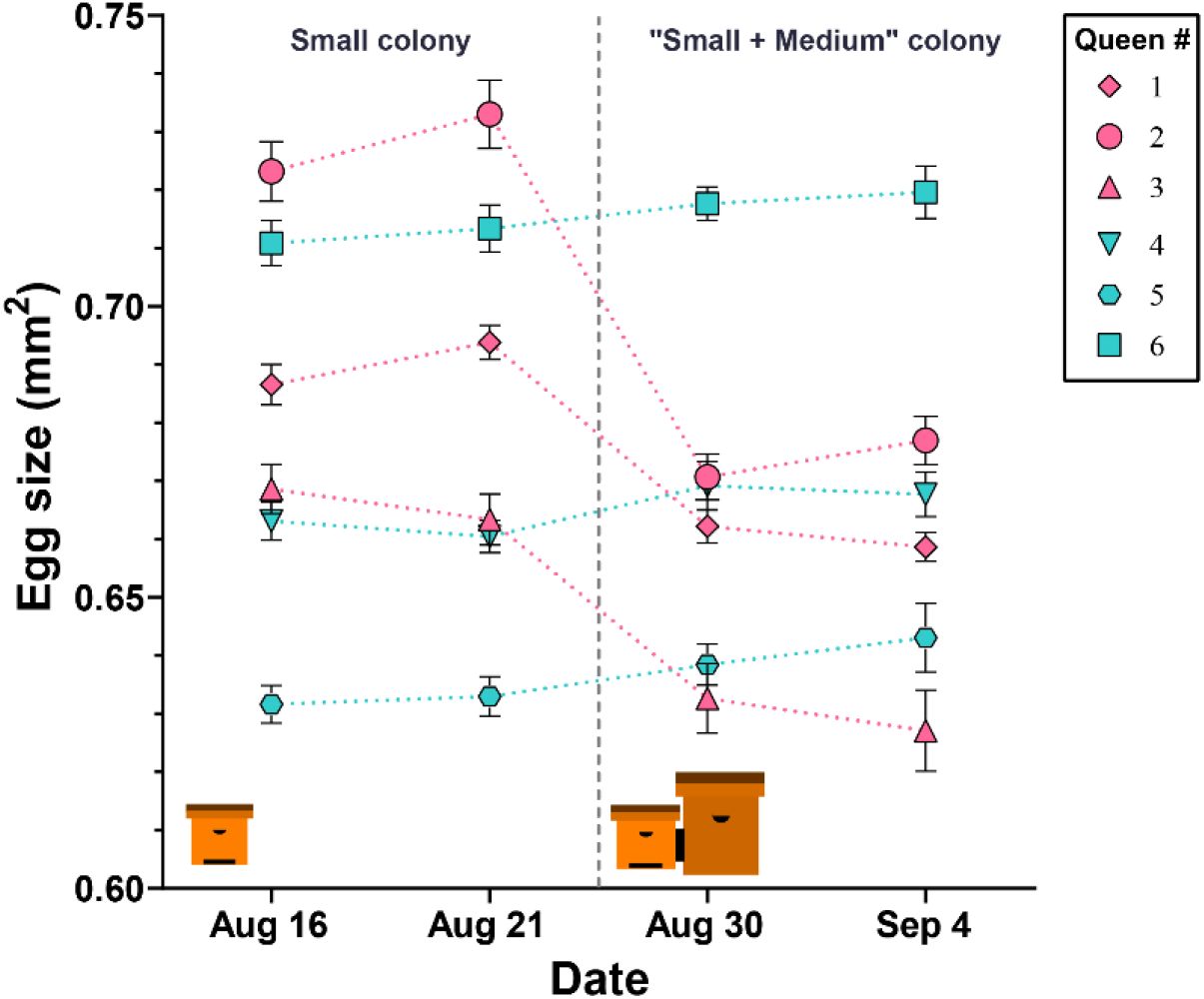
Egg size is actively regulated by the queen in response to perceived colony size. After initial egg size determination, queens in “Small” hives were either paired with an empty “Medium” hive box (controls: cyan color) or with a “Medium” hive box containing a colony (pink color). Queens in hives that were paired with another colony, decreased their egg size, while queens in control colonies maintained their egg sizes. Individual means ± S.D. are shown.

### Ovary proteome indicates that egg size is increased by up-regulation of cellular transport and metabolism

Comparing the ovary proteome of queens producing large eggs with that of queens producing small eggs identified a total of 2022 proteins. Among the 290 differentially expressed proteins, significantly more proteins were up-regulated (275) than down-regulated (15) in large egg-producing ovaries compared to small egg-producing ovaries (*χ*^2^ = 233.1, *p* < 0.001; Fig. 5*A* and Table S5).

**Fig. 5.**
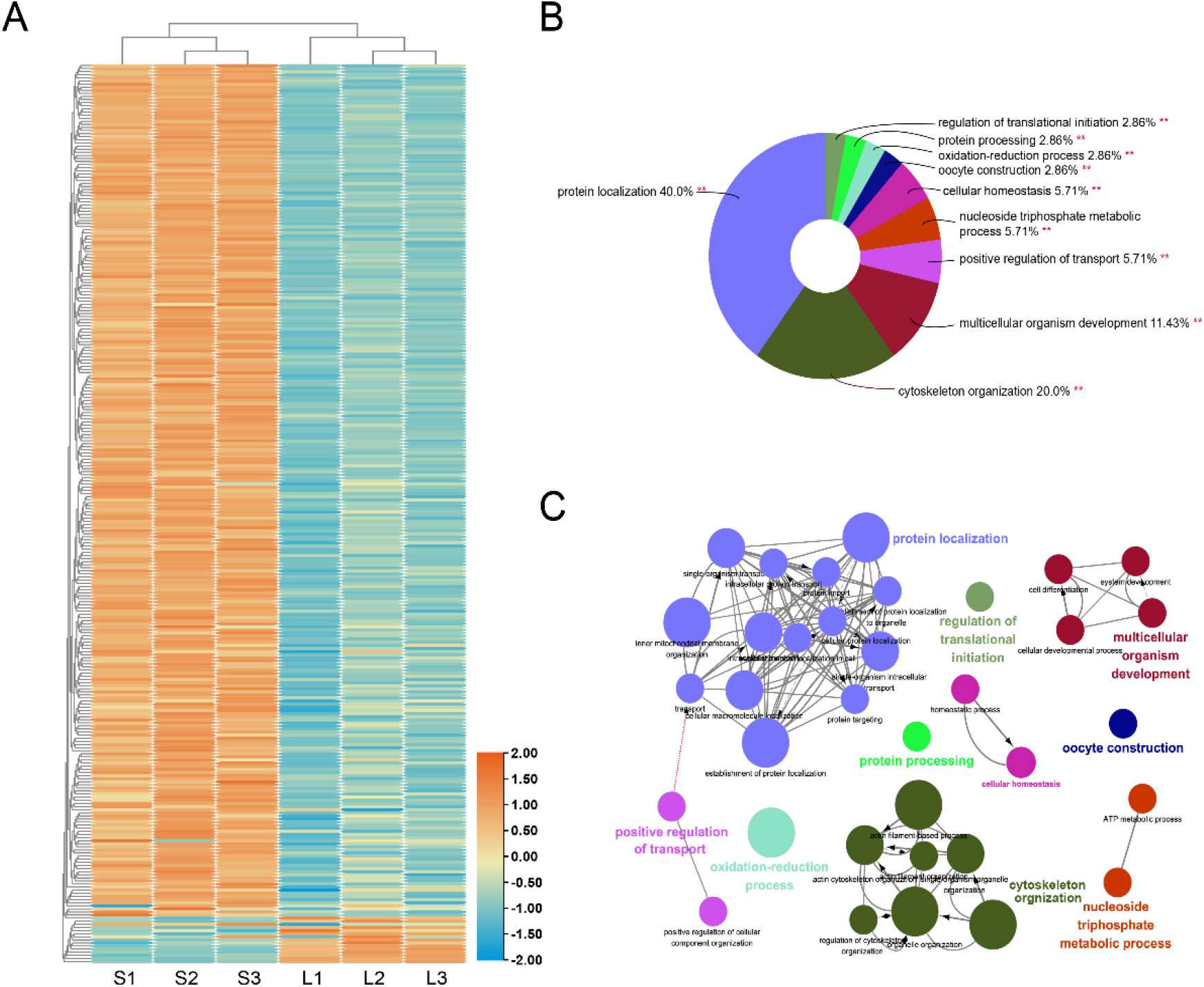
Quantitative protein differences between the ovaries of queens producing small and large eggs. The abundance of approximately 10% of all identified proteins was significantly different, with the vast majority of differences indicating up-regulation in the ovaries of queens that produced larger eggs because they were housed in small instead of large colonies **(A)**. Among the GO terms that were significantly (*p* < 0.01) enriched in the differentially abundant proteins, “protein localization” and “cytoskeleton organization” were most prominent **(B)**. Functional grouping of these overall GO terms, using kappa ≥ 0.4 as linking criterion confirmed that the GO terms represented at least 6 distinct functional groups **(C)**.

Gene ontology analysis of the proteomic data showed that the up-regulated proteins in large egg-producing ovaries s from queens in small colonies were significantly enriched in ten biological process terms (Fig. 5B and 5C and Table S6): “Protein localization” (*p* = 0.00004), “Oxidation-reduction process” (*p* = 0.00006), “Cytoskeleton organization” (p = 0.00009), “Cellular homeostasis” (*p* = 0.0003), “Protein processing” (*p* = 0.006), “Nucleoside triphosphate metabolic process” (*p* = 0.009), “Positive regulation of transport” (*p* = 0.010), “Multicellular organism development” (*p* = 0.011), “Oocyte construction” (*p* = 0.046), and “Regulation of translational initiation” (*p* = 0.048). In contrast, no GO term was significantly enriched in the down-regulated proteins.

The KEGG analysis revealed an enrichment of seven key pathways in the up-regulated proteins (Table S7), which included “Glycolysis / Gluconeogenesis” (*p* = 0.00002), “Citrate cycle (TCA cycle)” (*p* = 0.00003), “RNA transport” (*p* = 0.0003), “beta-Alanine metabolism” (*p* = 0.0005), “Protein processing in endoplasmic reticulum” (*p* = 0.0005), “Proteasome” (*p* = 0.0008), and “Oxidative phosphorylation” (*p* = 0.004). Consistent with the GO analysis, no enrichment could be identified in the proteins that were down-regulated in large egg-producing ovaries compared to small egg-producing ovaries.

The two largest GO term categories were “protein localization” and “cytoskeleton organization”. Of the 34 differentially expressed proteins that were associated with “cytoskeleton organization”, 29 were connected by a protein-protein interaction analysis (Fig. 6 and Table S8). This analysis pointed to five proteins with >10 connections to other proteins: Act5C (15), Rho1 (13), chic (12), Rac1 (11), and Tm2 (11). Instead of Act5C, the most connected protein with essential structural functions ^29^, we decided to further investigate the role of the second most connected protein Rho1, which represents a key regulator of cytoskeletal organization ^30^.

**Fig. 6.**
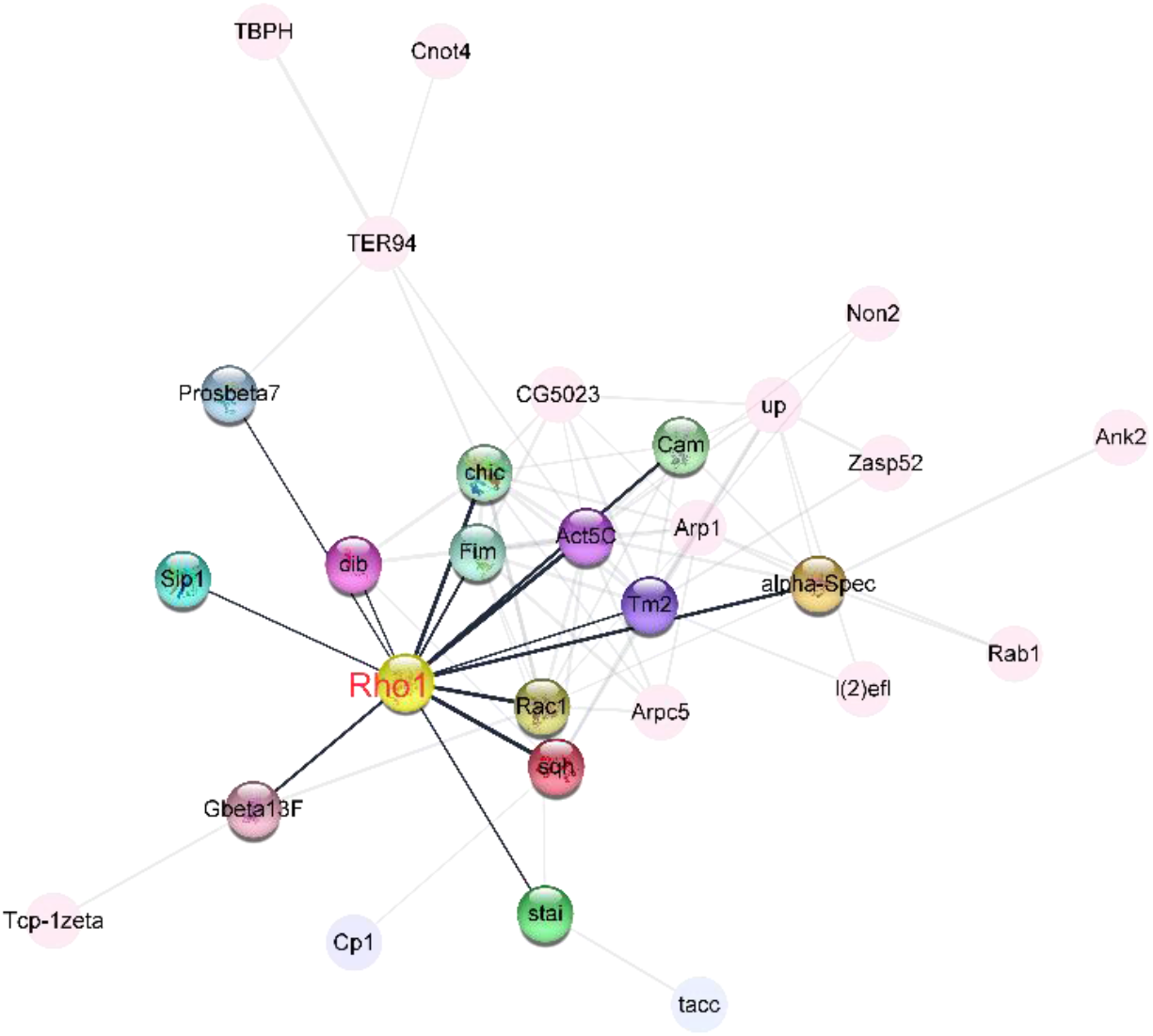
Central role of Rho1 in protein−protein interaction network of up-regulated cytoskeleton organization in large egg-producing honey bee queen ovaries. The interaction analysis, carried out in STRING v10), linked 29 proteins into the network. The highlighted nodes depict proteins that have a direct interaction with Rho1, a central regulator of cytoskeletal organization and the second most connected protein in the network.

### Rho1 in ovaries plays an important role in egg size regulation

Based on our proteomics results and functional evaluation of the top candidate genes, we hypothesized that *Rho1* is important for egg-size regulation. RNAscope® *in situ* hybridization enabled a fine-scale characterization of *Rho1* expression in the ovary, which was consistent with this hypothesis; little *Rho1* was expressed in the terminal filament but some expression was discernable in the germarium, concentrated in the cytocyst (incipient oocyte). Relative strong expression of *Rho1* was found in the growing oocytes of the vitellarium in contrast to nurse and follicle cells at that developmental stage. In mature oocytes, *Rho1* expression was again low (Fig. 7A). In the oocytes, *Rho1* was mainly located near the lateral cell cortex, which may represent areas of longitudinal growth (Fig. 7B).

**Fig. 7.**
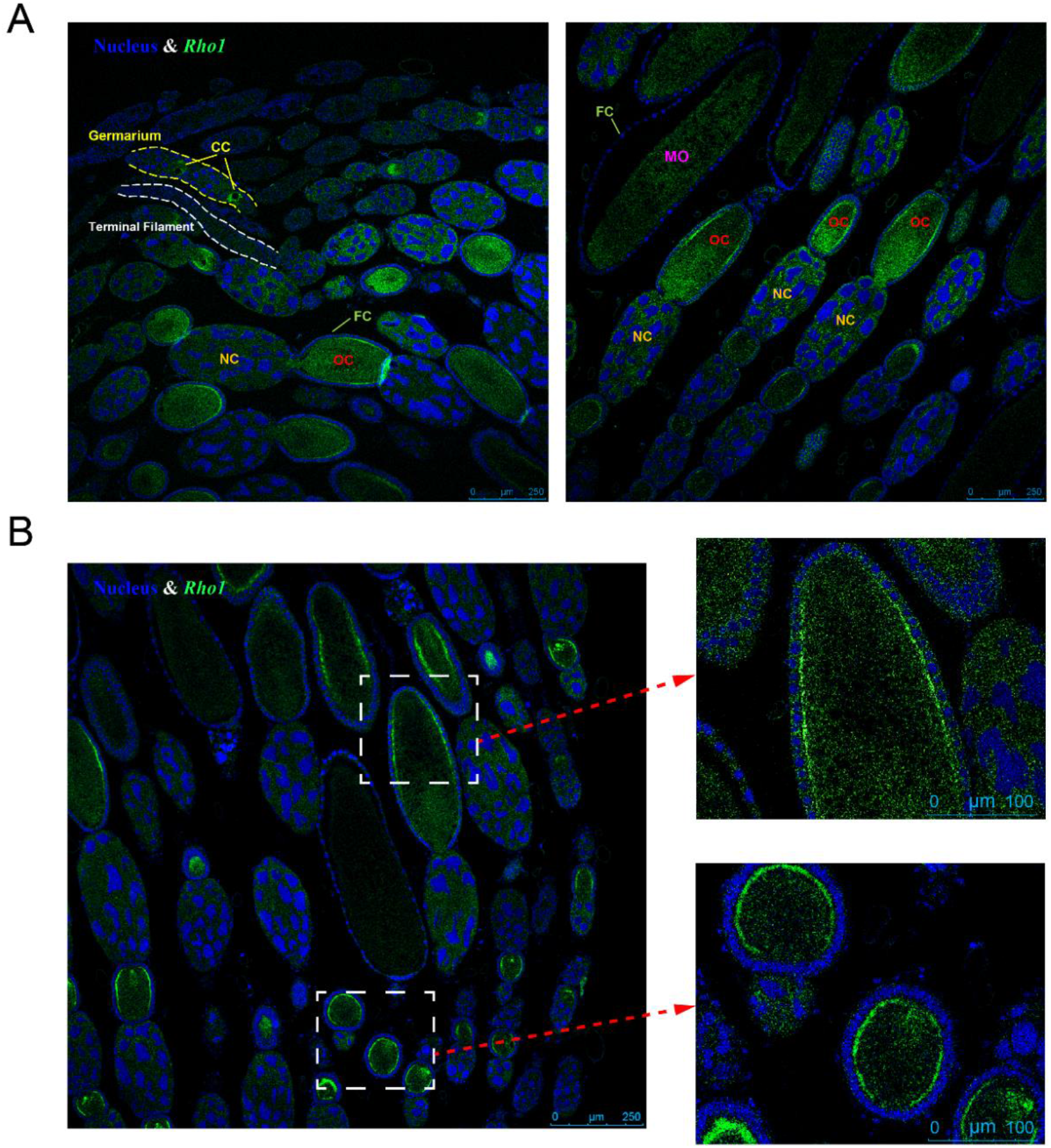
*Rho1* gene expression localization in the queen ovary via RNAscope® *in situ* hybridization. **(A)** The expression of *Rho1* (green color) is limited to the growth stages of the oocyte: In the germarium, *Rho1* is expressed in cytocysts (CCs) and in the vitellarium, *Rho1* is highly expressed in oocytes (OCs), particularly near the lateral cell surface **(B)**. In contrast, nurse cells (NCs) and follicle cells (FCs) do not exhibit elevated *Rho1* expression at this stage. Less expression of *Rho1* is observed in mature oocytes (MOs). Blue DAPI staining indicates cell nuclei for comparison.

RNAi-mediated knock-down of *Rho1* resulted in an average of 35.1% reduced *Rho1* expression compared to controls (Fig. 8A and Table S9). Expression of *Rho1* was also on average 57.0% higher in control queens from small colonies that produce large eggs than queens from large colonies that produce small eggs (Fig. 8A). The knock-down of *Rho1* consistently decreased egg sizes (Fig. 8*B* and Table S10) in all three queens in small colonies (Q10: F_(1,38)_ = 177.8, *p* < 0.001; Q11: F_(1,38)_ = 139.7, *p* < 0.001; Q12: F_(1,38)_ = 44.6, *p* < 0.001) and large colonies (Q4: F_(1,38)_ = 63.7, *p* < 0.001; Q5: F_(1,38)_ = 42.8, *p* < 0.001; Q6: F_(1,38)_ = 28.1, *p* < 0.001), while none of the six corresponding control queens exhibited significant egg size changes (F_(1,38)_ = 0.05 to 2.8, all *p* > 0.1). Thus, *Rho1* knock-down consistently reduced egg size even after the experimental queens increased (Q7-Q12 after transfer into small colonies: F_(1,38)_ = 45.6 to 654.8, all *p* < 0.001) or decreased (Q1-Q6 after transfer into large colonies: F_(1,38)_ = 24.8 to 158.4, all *p* < 0.001) the egg size that they had produced in medium-sized colonies at the start of the experiment (Fig. 8*B* and Table S10). All eggs appeared to be viable and differed phenotypically only in size. Across individuals from all treatment groups, *Rho1* expression at the end of the experiment correlated almost perfectly with the produced egg size (R_P_ = 0.98, n = 12, *p* < 0.001). The correlation between *Rho1* expression and egg size was confirmed in a second dataset of 12 queens that produced small and large eggs due to different colony sizes (R_P_ = 0.90, n = 12, *p* < 0.001).

**Fig. 8.**
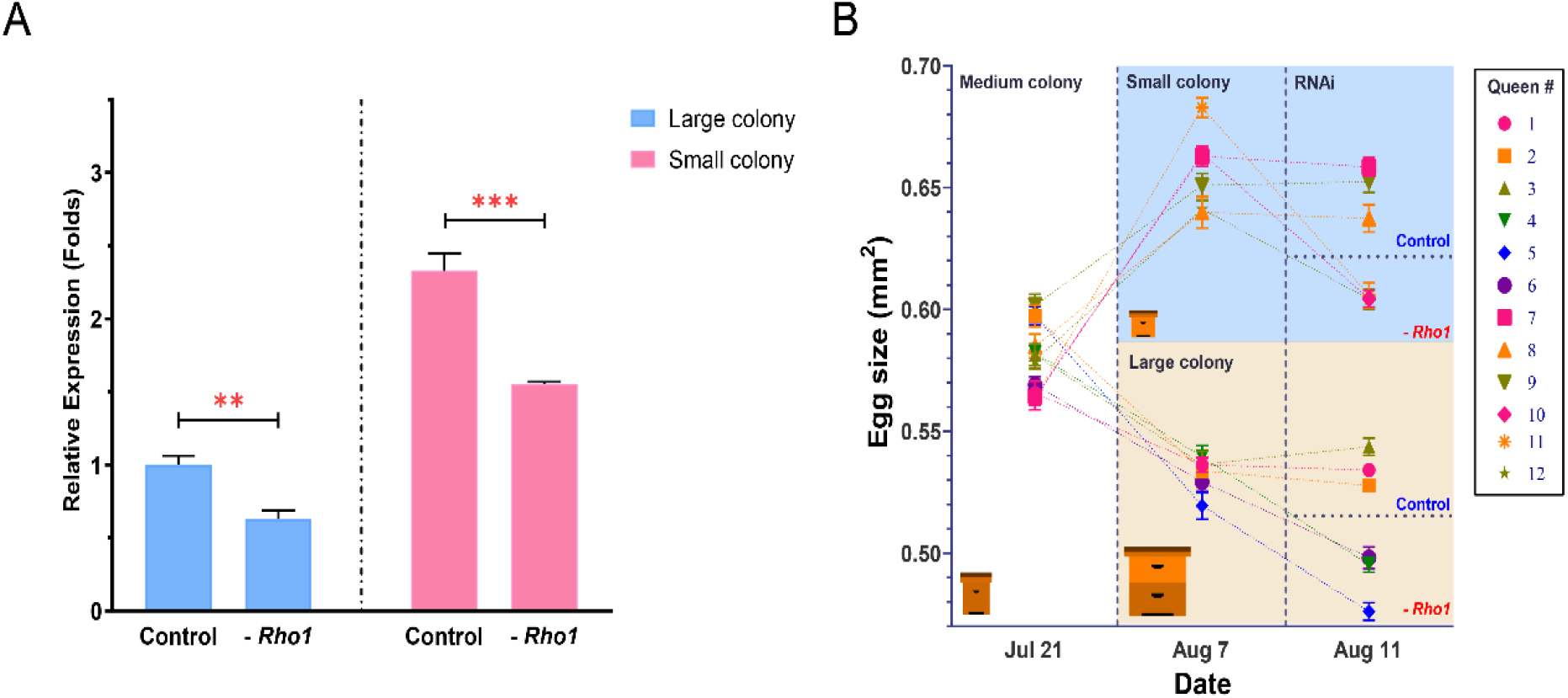
RNAi-mediated down-regulation of *Rho1* decreases egg size in both “Small” and “Large” colonies. **(A)** RT-qPCR results confirmed the experimental down-regulation of *Rho1* in ovaries of RNAi-injected queens and also showed that *Rho1* was significantly more expressed in queens that were housed in small colonies and thus produced larger eggs. **(B)** In this experiment, twelve sister queens were mated and introduced to medium-sized colonies to establish egg-laying. Subsequently, queens were randomly divided into two groups that were either introduced to small or large colonies. After the predicted egg-size differences were confirmed, three randomly-chosen queens in each group were injected with *Rho1*-siRNA mix, and the other three were injected with scramble siRNA. Final egg size measurements three days after injection demonstrated a significant reduction of egg size in all *Rho1* knockdown queens but not in control queens, regardless of colony environment.

## Discussion

The egg is the major physical connection between generations and thus central to inter-generational epigenetic effects that have major implications for offspring phenotypes ^31,32^ and life history evolution ^9,33^. Despite its importance and its considerable inter- and intra-specific variability, the egg life history stage remains poorly studied. Here, we provide evidence that egg size—a quantitative measure of maternal provisioning—is actively adjusted by honey bee queens in response to cues that relate to colony size. We also show that queens in smaller colonies have smaller ovaries but produce larger eggs. We find that protein localization, cytoskeleton organization, and energy generation are key proteomic changes in the ovary that mediate the production of large eggs. Finally, we identify the cytoskeleton organizer *Rho1* as a key regulator of the active egg size adjustment of honey bee queens.

Egg-size variation has been linked to parental or environmental conditions in numerous species ^6,10^, and we have previously provided evidence that honey bee queens also predictably adjust the size of produced eggs ^25^. The direction of egg-size adjustments is consistent between solitary species and honey bees; egg size is typically increased under unpredictable or unfavorable conditions ^34,35^ and positively correlated to maternal condition ^7^. We show here that these observations extend to honey bees as egg size is increased by the queen upon perception of a small colony. Small colonies may select for increased individual survival because each individual is proportionally more significant to the colony ^36^, but large egg size in small colonies could also be directly related to less consistent brood care and overall colony-level resource availability. Although we do not know how feeding rates of queens are affected by colony size, maternal condition may also influence egg size in honey bees. For example, older queens produce smaller eggs than young queens ^26^. However, our finding that queens in small colonies with reduced ovary size produce larger eggs suggests that a negative relationship at the individual level can also exist in honey bees. Such a negative relationship between maternal condition and egg size may also exist in other social insects where colony-level resource availability influences maternal resources and brood care performed by the workers ^37^.

The conventional trade-off between egg size and number ^5,6^ may not apply to honey bees because resources can be distributed from other colony members to the queen ^23,37,38^. Accordingly, little evidence for a trade-off between egg size and number was found in a previous study of honey bee queens ^25^. We demonstrate here that the restriction of egg laying by queens in large colonies does not lead to an increase in egg size, which is a prediction of the trade-off hypothesis. Instead, the observed egg size differences between queens in large and small colonies persist when egg laying rates are similar. Thus, the egg size differences represent active regulation instead of a passive consequence of egg laying rate. Active regulation is also compatible with egg-size adjustments in other contexts ^27^ and can explain why queens in food-restricted colonies also produce larger eggs ^25^. Nevertheless, queens in large colonies typically produce more eggs than queens in small colonies. Our finding that queens in large colonies have heavier ovaries indicates a physiological adaptation to satisfy the egg-laying demand in large colonies ^39^, but our phenotypic and molecular results indicate that such physiological limitations are not the cause of the observed egg size plasticity.

Our study further demonstrates that direct resource availability cannot explain the differences in egg size produced by honey bee queens in small versus large colonies. We find that connecting small colonies to another, larger colony without any physical contact leads to a reduction in egg size that is similar to the effect seen when queens are transferred between these colony types. Thus, our results suggest that the perception of colony size by the queen is sufficient for her to adjust the size of her eggs. We can exclude direct physical contact among individuals, which is used by worker honey bees to assess colony size ^40^, and thus we report a new modality by which honey bees perceive colony size. Multiple cues that travel through a double-screened tunnel could prompt the egg size adjustment in queens, including sound, temperature, or pheromones and other semiochemicals. Future distinction among these possibilities will permit a subsequent investigation of the mechanisms by which social cues are translated into the physiological adjustments inside the ovary that we document here.

Broad comparisons find pronounced influences of social structure and behavior on egg size because variation in parental care is intricately linked to the initial investment in eggs ^41,42^. Much less is known about social factors that lead to individual egg-size plasticity ^43^, particularly in cases that are as dynamic and reversible as illustrate here. We have not attempted to measure the speed at which queens adjust their egg size, but it could be much faster than the 1-2 weeks provided to queens in our experiments. Adjustments may even be made instantaneously, as queen- and worker-destined eggs differ in size ^27^ even though they are presumably produced at almost the same time. The advantageous caste bias of larger eggs ^28^ should select for paternal effects to increase egg size, in contrast to other polyandrous species in which males predominantly manipulate female fecundity ^44^. Egg-size variation due to paternal manipulation remains to be investigated in honey bees in the context of the strong maternal control over egg size that we demonstrate in this study.

Due to the general paucity of *a priori* information on molecular mechanisms that determine egg size in insects ^14^, we used a naïve, quantitative proteomic comparison to identify the molecular causes of the egg-size plasticity in honey bee queens. The quantity of numerous proteins is associated with the production of either large or small eggs. Plod, which controls egg length in *Drosophila* ^45^, is not among these proteins, but collagen IV, which may influence egg size in *Drosophila* indirectly ^46^, is found more abundantly in large egg-producing ovaries. The vast majority (almost 95%) of differently abundant proteins are up-regulated in ovaries that produce large eggs. Thus, the anatomically smaller ovaries are physiologically more active in several key processes than the larger ovaries that produce smaller eggs. The GO-enrichment analysis indicated that the two largest up-regulated processes are “protein localization” and “cytoskeletal regulation”, while several energy metabolic processes are highlighted by the KEGG-pathway analysis. These functional categories indicate that egg-size variation is not a simple increase of egg volume but reflects real differences in offspring provisioning, although the proteome of small and large eggs remains to be characterized ^47^. Higher energy generation may be needed to produce more costly large eggs ^48^, and the cytoskeleton and protein localization processes are key to loading the egg with nutrients in polytrophic ovaries ^49,50^. Several of the other GO terms, such as “multicellular organism development” and “oocyte construction”, are further plausible candidates to explain some of the observed variation in egg size.

Among all involved processes, we considered “cytoskeletal organization” as the most likely regulatory mechanism, while other processes are more likely involved in downstream effector functions. Thus, we identified Rho1 as a potential candidate because it was centrally connected in the protein-protein interaction network of up-regulated proteins related to cytoskeletal organization and a plausible functional candidate: *Rho1* is a small, conserved GTPase with a likely role in egg size regulation ^51^. *Rho1* has multiple functions but generally plays an important role in cell morphogenesis by regulating the cytoskeleton ^52^. It primarily has been implicated in actin regulation ^52^, which is itself important for insect oogenesis ^53^ but can also indirectly influence the microtubule network ^54^. The regulation of *Rho1* activity is complex ^55^ and multiple participating signaling pathways could transduce extracellular signals in the ovary into cytoskeletal reorganization of the eggs.

The observed spatio-temporal expression of *Rho1* observed in our RNAScope® experiment conforms well with the hypothesis that *Rho1* influences egg growth in the vitellarium. The genetics of honey bee eggs is not yet well-developed ^50,56^ and early developmental studies in general are mostly focused on pattern formation. We identify the spatio-temporal expression pattern of *Rho1* and some likely interaction partners, but the molecular function of *Rho1* remains to be elucidated. However, the role of *Rho1* in egg-size determination is further supported by the consistent decrease of egg size when *Rho1* is knocked-down via siRNA injection. This specific effect occurs robustly in small- and large-egg producing queens. Resulting eggs only differed in size, suggesting that the knock-down of *Rho1* does not cause pathological effects. The practical difficulties of genetic engineering in honey bees ^57^ prohibited a complementary gain-of-function experiment. The almost perfect correlation between *Rho1* expression and egg size of queens across two different colony sizes and RNAi treatment groups strengthens the interpretation that social conditions, inter-individual differences and RNAi manipulation all act through *Rho1* in a comparable manner to influence egg size. This conclusion was supported by a tight correlation between egg size and *Rho1* expression in a second, independent data set.

Honey bee queens also adjust their egg size depending on whether a worker- or queen-destined egg is laid ^27^, and egg size influences the probability that an egg is raised into a future queen ^28^ and her reproductive quality ^27^. Our comparative results of the ovary proteome suggest that larger eggs are indeed of superior quality, but a comparison of the actual egg content remains to be performed to substantiate this argument. It is likely but also remains to be tested whether *Rho1* causes egg-size variation in honey bees in other contexts, such as maternal genotype ^25^ or age ^26^, and could be used as a honey bee health indicator. In bumblebees, egg size is negatively impacted by pesticide stress ^58^. More generally, our findings also suggests that this conserved gene may regulate egg size in other species.

## Methods

### Experimental model and subject details

All studies were conducted in the Western honey bee, *Apis mellifera*, using colonies of mixed origin and derived from commercial populations, that were kept in the research apiary of the University of North Carolina at Greensboro, NC, USA (UNCG: 2020) or in the research apiary of the Institute of Apicultural Research in Beijing, China (IAR: 2021). We used standard husbandry methods to house experimental colonies ^59^, monitoring and adjusting colony size and food status but refraining from any other treatments during the experiments. We defined three distinct colony sizes: “Small” colonies contained 500–700 worker bees housed in mating hives (nucs) equipped with three half-frames of medium depth, “Medium” colonies contained 2,500–3,500 workers bees housed in a 5-frame Langstroth hive box with standard frames, and “Large” colonies with 8,000–9,000 worker bees in a standard 8-frame Langstroth hive. Each separate experiment was conducted with a set of sister queens that we raised using standard methods ^59^ and allowed to mate naturally.

### Repeated transfer experiments

As an extension of our previous study ^25^, an experiment was set up in the UNCG apiary to transfer one group of queens from “Medium” to “Small” to “Large” colonies and simultaneously transfer another group from “Medium” to “Large” to “Small” colonies. During each stage, egg size was measured from 20 eggs per queen twice (one week apart). The measurements followed our previous protocol ^25^, where eggs produced overnight were randomly selected in the next morning and transferred with a grafting tool from standard worker brood cells onto a 0.01 mm stage micrometer (Olympus, Japan). Eggs were laterally photographed under threefold magnification while ensuring that the egg was completely level. Each photo was then processed with the open-source ImageJ software (version 1.52p; National Institutes of Health, USA) by manually tracing the egg’s outline using the polygon-selection tool. The selected area was measured in mm^2^ (note that in our previous work ^25^, a simple conversion mistake led to erroneous μm^2^ units) as a representation of egg size.

From an original 16 queens, 11 were successfully mated and started egg-laying in their respective “Medium” colonies. After two weeks and two egg size measurements (on the 12^th^ and 19^th^ of June 2020), six of these queens were transferred to “Small” colonies, while the remaining five were transferred to “Large” colonies. After two weeks of acclimation, another two egg-size measurements from each queen were performed (on the 7^th^ and 14^th^ of July 2020) before reciprocally transferring queens between “Small” and “Large” colonies. This transfer was survived by five queens ending up in “Small” colonies and three queens in “Large” colonies. After another two weeks of acclimation, egg size of these remaining queens was measured as before (on the 4^th^ and 11^th^ of August 2020).

At the conclusion of this experiment, all surviving queens were weighed and sacrificed for determination of their body size, ovary weight, and ovary proteome. To increase sample size, two additional queens were transferred from “Medium” colonies and housed in “Small” colonies for two weeks before including them in these analyses. Queens were captured alive and weighed in a pre-weighed 1.5 ml tube centrifuge tube to the nearest microgram. After cold-anaesthetization, both forewings were detached from the queens and mounted on a microscope slide to determine the distance between the distal end of the marginal cell and the intersection of the Cu1 and 2m-cu veins as a representative size measure ^60^. The values from both wings were averaged. Subsequently, the ovary was dissected from the chilled abdomen, weighed, and frozen at -80°C for proteome profiling (see below).

A simpler, additional experiment was conducted to perform another comparison of queen- and ovary weight between queens in “Small” and “Large” colonies in order to gather more ovaries from these treatment groups for proteome profiling. Accordingly, sister queens were reared from a randomly selected mother in the UNCG apiary. After their maturation into ovipositing queens in “Medium” colonies, four were successfully introduced into “Small” colonies and four others into “Large” colonies. After two weeks, the production of large and small eggs respectively was confirmed for all eight accepted queens (Table S2) and queens were compared with regard to their body weight, body size, and ovary weight. The ovaries of these queen were also collected and stored at -80°C for further use.

A third study of ovary size was performed with all queens at the end of the RNAi knock-down experiment (see below). For this purpose, the body weight, the wing size, and the ovary weight of the 12 queens were measured as described above. In this instance, the body weight was measured before injection, while the wing size and the ovary weight were measured after injection.

### Oviposition restriction experiment

Although no significant correlation between egg size and number was found in our previous experiments ^25^, we tested the hypothesis that egg size is a passive consequence of different egg-laying rates more explicitly. Eight sister queens were reared from a randomly selected source in the IAR apiary in July 2021, introduced as mature queen cells to “Medium” colonies for emergence, mating, and initiation of oviposition. Subsequently, the queens were introduced into “Large” colonies, and two weeks after acceptance their egg sizes were measured as described above. Subsequently, queens were randomly split into an oviposition restriction group and an unmanipulated control group. Oviposition restriction was achieved by caging queens for two weeks in their hives on top of capped brood combs without empty cells as egg-laying opportunities. Immediately after these two weeks, all queens were caged on identical sections of comb with empty cells to measure their egg sizes again. Each queen was evaluated separately for significant differences in egg size between the start and end of the experiment.

### Colony extension experiment

To clarify how colony size influences queen oviposition, we tested whether physical contact or material transfers are necessary to alter the size of eggs produced by the queen. Six sister queens were reared from a randomly selected mother in the IAR apiary in July 2021. After maturation (as described above), these queens were introduced into “Small” colonies. After an acclimation period of two weeks, egg sizes produced by all queens were determined twice as described above (15^th^ and 21^st^ of August 2021). The colonies were then connected via a double-screened tunnel to a “Medium” hive that contained either empty comb (control) or a “Medium” colony with corresponding amounts of food, brood, and workers, but no queen (treatment). Tunnels were 3 cm long, 10 cm wide, and 20 cm high. Both ends were screened with fine wire mesh to prevent any physical contact among the workers in opposing hives. Worker drifting between hives was prevented by pointing the hive entrances of the two connected units to opposite directions, as well as coloring and designing the entrances differently. One week later, egg-size measurements were performed (30^th^ of August) and repeated once after an additional week (4^th^ of September).

### Ovary proteome analysis

In an unbiased search for differences, the protein content of ovaries that produce small eggs (from queens in “Large” colonies) and ovaries that produce large eggs (from queens in “Small” colonies) was studied with a label-free LC-MS/MS approach. A total of 18 ovaries, collected during the two UNCG experiments described above, were included. For both groups (small egg-producing queens from large colonies and large egg-producing queens from small colonies), nine ovaries were pooled randomly into three biological replicates.

Total protein was extracted using previously described methods ^61^. Protein concentration was determined using a Bradford assay and the general quality of extracted proteins was confirmed by SDS-PAGE with Coomassie Blue staining. An aliquot of 200 μg of protein from each pool was reduced with DTT (final concentration 10 mM) for 1 h, then alkalized with iodoacetamide (final concentration 50 mM) for 1 h in the dark. Thereafter, protein samples were digested at 37°C overnight with sequencing grade trypsin (enzyme: protein (w/w) = 1:50). The digestion was stopped by adding 1μl of formic acid then desalted using C18 columns (Thermo Fisher Scientific, USA). The desalted peptide samples were dried and dissolved in 0.1% formic acid in distilled water, then quantified using a Nanodrop 2000 spectrophotometer (Thermo Fisher Scientific) and stored at -80°C for subsequent LC-MS/MS analysis.

LC-MS/MS analysis was performed on an Easy-nLC 1200 (Thermo Fisher Scientific) coupled Q-Exactive HF mass spectrometer (Thermo Fisher Scientific). Buffer A (0.1% formic acid/water) and buffer B (0.1% formic acid and 80% acetonitrile in water) were used as mobile phase buffers. Peptides were separated using a reversed-phase trap column (2 cm long, 100 μm inner diameter, filled with 5.0 μm Aqua C18 beads; Thermo Fisher Scientific) and an analytical column (15 cm long, 75 μm inner diameter, filled with 3 μm Aqua C18 beads; Thermo Fisher Scientific) at a flow rate of 350 nL/min with the following 120 min gradients: from 3 to 8% buffer B in 5 min, from 8 to 20 % buffer B in 80 min, from 20 to 30% buffer B in 20 min, from 30 to 90% buffer B in 5 min, and remaining at 90% buffer B for 10 min. The eluted peptides were injected into the mass spectrometer via a nano-ESI source (Thermo Fisher Scientific). Ion signals were collected in a data-dependent mode and run with the following settings: scan range: m/z 300-1,800; full scan resolution: 70,000; AGC target: 3E6; MIT: 20 ms. For MS/MS mode, the following settings were used. Scan resolution: 17,500; AGC target: 1E5; MIT: 60 ms; isolation window: 2 m/z; normalized collision energy: 27; loop count 10; dynamic exclusion: 30 s; dynamic exclusion with a repeated count: 1; charge exclusion: unassigned, 1, 8, >8; peptide match: preferred; exclude isotopes: on. The corresponding raw data were retrieved using Xcalibur 3.0 software (Thermo Fisher Scientific).

The extracted MS/MS spectra were searched against the protein database of *Apis mellifera* (23,430 sequences, from NCBI) appended with the common repository of adventitious proteins (cRAP, 115 sequences, from The Global Proteome Machine Organization) using PEAKS 8.5 software (Bioinformatics Solutions, Canada). The search parameters were: ion mass tolerance, 20.0 ppm using monoisotopic mass; fragment ion mass tolerance, 0.05 Da; enzyme, trypsin; allow non-specific cleavage at none end of the peptide; maximum missed cleavages per peptide, 2; fixed modification, Carbamidomethylation (C, +57.02); variable modifications, Oxidation (M, +15.99); maximum allowed variable PTM per peptide, 3. A fusion target-decoy approach was used for the estimation of false discovery rate (FDR) and controlled at ≤ 1.0% (−10 log P ≥ 20.0) both at peptide and protein levels. Proteins were identified based on at least one unique peptide.

Quantitative comparison of the egg proteome between the two experimental groups was performed by the label-free approach embedded in PEAKS Q module. Feature detection was performed separately on each sample by using the expectation-maximization algorithm. The features of the same peptide from different samples were reliably aligned together using a high-performance retention time alignment algorithm. Significance was calculated by ANOVA, using a threshold of *p* ≤ 0.01). Results were visualized as a heatmap using TBtools software ^62^, clustering based on Euclidean distance and the “complete” method.

For further functional analysis, honey bee proteins were mapped to their *Drosophila melanogaster* homologs using KOBAS 3.0 ^63^. Proteins of interest were uploaded as fasta sequences, *Drosophila melanogaster* was selected as target species, and similarity mapping was conducted with default cutoffs (BLAST *E*-value < 1E−5 and rank ≤ 5).

Functional Gene Ontology (GO) enrichment analysis of the quantitatively different proteins was performed based on biological processes with ClueGO + CluePedia version 2.5.7 ^64^, a Cytoscape (version 3.8.2) plugin. Two-sided hypergeometric test (enrichment/depletion) with *p*-value ≤ 0.05 was used followed by Bonferroni correction. The GO tree interval was set between 3 and 8, with minimum 5 genes and 1% of genes. Kappa score ≥ 0.4 was applied to generate term-term interrelations and functional groups based on shared genes between the terms. KEGG pathway enrichment analysis was done in Metascape (http://metascape.org/) with default settings: minimum overlap, 3; *p*-value cutoff, 0.01; minimum enrichment, 1.5.

For the exploration of functional protein connections involved in the major enriched biological process terms, protein−protein interaction (PPI) networks were constructed among the differing proteins in STRING ^65^. A full STRING network was built with medium confidence (0.4) and FDR < 5%. The PPI networks were visualized using Cytoscape (version 3.8.2).

### Examination of expression patterns of *Rho1*

Based on the proteomic analyses, the small GTPase *Rho1* emerged as a candidate regulator of egg size during honey bee oogenesis, which motivated us to study its expression patterns in the ovary and inside the oocyte by RNAscope® *in-situ* hybridization ^66^. The probes were designed and prepared by Advanced Cell Diagnostics (ACD, Inc., Hayward, USA) and an RNAscope® Fluorescent Multiplex Reagent kit (ACD) was used following the manufacturer’s instructions. Immediately following dissection, the ovary tissues of randomly selected, mature queens from the IAR apiary were fixed in 10% neutral buffered formalin for 32 h at room temperature (RT). Thereafter, the samples were dehydrated using a standard ethanol series, followed by xylene. The dehydrated samples were embedded in paraffin and then cut into 1 μm sections using a RM2235 microtome (Leica, Germany) that were gently deposited onto glass microscope slides. The slides were then baked for 1 h at 60°C and deparaffinized at RT. The sections were treated with hydrogen peroxide for 10 min at RT and then washed with fresh distilled water.

Subsequently, the target retrieval step was performed using 1×RNAscope® target retrieval reagent. The slides were air dried briefly and then boundaries were drawn around each section using a hydrophobic pen (ImmEdge® pen; Vector Laboratories, USA). After hydrophobic boundaries had dried, the sections were incubated in protease IV reagent for 2 min, followed by a 1×PBS wash. Each slide was then placed in a prewarmed humidity control tray (ACD) containing dampened filter paper and incubated in a mixture of Channel 1 probes (*Rho1*, ACD catalog #1061331-C1) for 2 h in the HybEZ® oven (ACD) at 40°C. Following probe incubation, the slides were washed two times in 1×RNAscope® wash buffer and returned to the oven for 30 min after submersion in AMP-1 reagent. Washes and hybridization were repeated using AMP-2, AMP-3, and HRP-C1 reagents with a 30 min, 15 min, and 15 min incubation period, respectively. The slides were then submerged in TSA® Plus FITC and returned to the oven for 30 min. After washing two times in 1×RNAscope® wash buffer, the slides were incubated with HRP blocker for 15 min in the oven at 40°C. Finally, the slides were washed two times in 1×RNAscope® wash buffer and incubated with DAPI for 1 min. The images were visualized with a Leica SP8 (Leica) confocal microscope and acquired with the sequence program of the Leica LAS X software.

### RNAi-mediated down-regulation of Rho1

To test the hypothesis that *Rho1* expression controls the size of the eggs that honey bee queens produce, we investigated the effects of RNAi-mediated down-regulation of *Rho1*. Four specific siRNAs targeting Rho1 of *Apis mellifera* (GenBank: LOC409910) were designed and synthesized by GenePharma RNAi Company (Shanghai, China). Scrambled siRNA of random sequence was used as a negative control (GenePharma). For all siRNA sequences see Table S11.

Twelve sister queens were produced from a random source hives of the IAR apiary. They were introduced into “Medium” colonies to mate and establish egg laying. When a regular laying pattern was established, the size of the eggs produced by all queens was measured as described above. Then, queens were randomly divided into two groups of six that were either introduced into “Small” or “Large” colonies. Two weeks after queen acceptance, egg size measurements were repeated. One day later, three queens in each group were randomly selected and injected with 1 μl/queen of *Rho1*-siRNA mix (mixture of the four *Rho1*-siRNAs, 1 μg/μl), and the other three were injected with 1 μl/queen of scrambled siRNA (1 μg/μl). The queens were transferred to the laboratory and narcotized with CO_2_ before injection. Injections were made dorsally between the 4th and 5th abdominal segment of queens using a microliter syringe (NanoFil®; World Precision Instruments, USA) coupled with a 35G needle (NF35BV-2; World Precision Instruments). Injected queens were given time to recover and placed back into their original colonies. Egg size measurements for each queen were performed three days after injection and control or treatment effects on egg size were tested separately for each queen by comparing the sizes produced before and after RNAi injection.

To assess the efficacy of RNAi knock-down of *Rho1* and investigate the correlation between ovary size and *Rho1* expression, the expression of *Rho1* was quantified by quantitative real-time PCR. Queens were anaesthetized before dissection of the ovary for weight measurement (see above) and subsequent RNA extraction, cDNA synthesis, and RT-qPCR according to previously described methods ^67^. The average of three technical replicates were computed and used in subsequent analyses. Reference genes were evaluated by GeNorm analysis ^68^ that indicated all evaluated reference genes (*Arfgap3, CylD, GAPDH, Keap1, Kto, mRPL44, RpA*-*70, Rpn2*) had high expression stability expression across samples (M-values < 0.4). Following GeNorm’s recommendation, we used *Arfgap3* and *CylD* to calculate relative gene expression as 2^-ΔΔCt ^69^. A corresponding analysis to confirm the correlation between *Rho1* and egg size was performed in a second set of twelve queens without RNAi exposure.

## Data availability

The LC−MS/MS data and search results were deposited in ProteomeXchange Consortium (http://proteomecentral.proteomexchange.org) via the iProX partner repository with the dataset identifier IPX0002748002 (https://www.iprox.cn/page/PSV023.html;?url=1639928825446wOdF, Password: iE8A).

## Acknowledgments

We thank all our lab members for their encouragement and support. This study was funded by grants from the National Natural Science Foundation of China (31970428) and the China Scholarship Council (201903250009) to B.H., a scholarship to E.A. by the US National Research Council, a grant to S.X. by the Agricultural Science and Technology Innovation Program (CAAS-ASTIP-2015-IAR), a grant to J.L. by the earmarked fund for Modern Agro-Industry Technology Research System (CARS-44) in China, and grants from the US Army Research Office (W911NF1920161) and the Alberta Beekeepers Commission to O.R.

## Author Contributions

B.H., E.A., S.X., and O.R. designed the research; B.H., Q.W., E. A., H.H. and L.M. performed the research; B.H., E.A. and O.R. analyzed the data and interpreted the results; O.R. J.L., and S.X. provided resources, B.H., E.A. and O.R. wrote the paper; and J.L., S.X., D.R.T. and M.K.S. edited the manuscript.

## Materials & Correspondence

Correspondence and requests for materials should be addressed to Olav Rueppell.

## Supplementary information

Table S1. Egg size measurements for first experiment.

Table S2. Various queen and egg size measurements from follow-up experiments.

Table S3. Egg size measurements from oviposition restriction experiment.

Table S4. Egg size measurements from colony connection experiment.

Table S5. Proteins of difference abundance.

Table S6. Gene Ontology results of upregulated proteins in large-egg producing ovaries.

Table S7. KEGG Pathway results of upregulated proteins in large-egg producing ovaries.

Table S8. Protein-protein interaction analysis of cytoskeleton organization proteins.

Table S9. Gene expression data from RNAi experiment.

Table S10. Egg size measurements from RNAi experiment.

Table S11. Sequence information for RNAi and qPCR.

## References

1. Stearns, S.C. (1989). Trade-offs in life-history evolution. Functional Ecology 3, 259–268.

2. Flatt, T. (2020). Life-history evolution and the genetics of fitness components in Drosophila melanogaster. Genetics 214, 3–48.

3. Bonsall Michael B., Jansen Vincent A. A., and Hassell Michael P. (2004). Life history trade-offs assemble ecological guilds. Science 306, 111–114.

4. Van Noordwijk, A.J., and de Jong, G. (1986). Acquisition and allocation of resources: their influence on variation in life history tactics. The American Naturalist 128, 137–142.

5. Smith, C.C., and Fretwell, S.D. (1974). The optimal balance between size and number of offspring. The American Naturalist 108, 499–506.

6. Dani, K.G.S., and Kodandaramaiah, U. (2017). Plant and animal reproductive strategies: Lessons from offspring size and number tradeoffs. Frontiers in Ecology and Evolution 5, 38.

7. Fox, C.W., and Czesak, M.E. (2000). Evolutionary ecology of progeny size in arthropods. Annual Review of Entomology 45, 341–369.

8. Berrigan, D. (1991). The allometry of egg size and number in insects. Oikos 60, 313–321.

9. Räsänen, K., and Kruuk, L.E.B. (2007). Maternal effects and evolution at ecological time-scales. Functional Ecology 21, 408–421.

10. Sánchez-Tójar, A., Lagisz, M., Moran, N.P., Nakagawa, S., Noble, D.W., and Reinhold, K. (2020). The jury is still out regarding the generality of adaptive ‘transgenerational’effects. Ecology Letters 23, 1715–1718.

11. Bebbington, K., and Groothuis, T.G.G. (2021). Who listens to mother? A whole-family perspective on the evolution of maternal hormone allocation. Biological Reviews 96, 1951–1968.

12. Christians, J.K. (2002). Avian egg size: variation within species and inflexibility within individuals. Biological Reviews 77, 1–26.

13. Azevedo, R.B., Partridge, L., and French, V. (1997). Life-history consequences of egg size in Drosophila melanogaster. The American Naturalist 150, 250–282.

14. Jha, A.R., Miles, C.M., Lippert, N.R., Brown, C.D., White, K.P., and Kreitman, M. (2015). Whole-genome resequencing of experimental populations reveals polygenic basis of egg-size variation in Drosophila melanogaster. Molecular Biology and Evolution 32, 2616–2632.

15. Abbot, P., Abe, J., Alcock, J., Alizon, S., Alpedrinha, J.A.C., Andersson, M., Andre, J.B., van Baalen, M., Balloux, F., Balshine, S., et al. (2011). Inclusive fitness theory and eusociality. Nature 471, E1–E4.

16. Boomsma, J.J., and Gawne, R. (2018). Superorganismality and caste differentiation as points of no return: how the major evolutionary transitions were lost in translation. Biological Reviews 93, 28–54.

17. Oster, G.F., and Wilson, E.O. (1978). Caste and Ecology in the Social Insects. (Princeton University Press).

18. Flatt, T., Amdam, G.V., Kirkwood, T.B., and Omholt, S.W. (2013). Life-history evolution and the polyphenic regulation of somatic maintenance and survival. The Quarterly Review of Biology 88, 185–218.

19. Lee, R.D. (2003). Rethinking the evolutionary theory of aging: Transfers, not births, shape social species. Proc Natl Acad Sci USA 100, 9637–9642.

20. Shik, J.Z., Hou, C., Kay, A., Kaspari, M., and Gillooly, J.F. (2012). Towards a general life-history model of the superorganism: predicting the survival, growth and reproduction of ant societies. Biology Letters 8, 1059–1062.

21. Negroni, M.A., Jongepier, E., Feldmeyer, B., Kramer, B.H., and Foitzik, S. (2016). Life history evolution in social insects: a female perspective. Current Opinion in Insect Science 16, 51–57.

22. Keller, L., and Genoud, M. (1997). Extraordinary lifespans in ants: a test of evolutionary theories of ageing. Nature 389, 958–960.

23. Schrempf, A., Giehr, J., Röhrl, R., Steigleder, S., and Heinze, J. (2017). Royal Darwinian demons: enforced changes in reproductive efforts do not affect the life expectancy of ant queens. The American Naturalist 189, 436–442.

24. Hall, J.M., Mitchell, T.S., Thawley, C.J., Stroud, J.T., and Warner, D.A. (2020). Adaptive seasonal shift towards investment in fewer, larger offspring: Evidence from field and laboratory studies. Journal of Animal Ecology 89, 1242–1253.

25. Amiri, E., Le, K., Melendez, C.V., Strand, M.K., Tarpy, D.R., and Rueppell, O. (2020). Egg-size plasticity in Apis mellifera: Honey bee queens alter egg size in response to both genetic and environmental factors. Journal of Evolutionary Biology 33, 534–543.

26. Al-Lawati, H., and Bienefeld, K. (2009). Maternal age effects on embryo mortality and juvenile development of offspring in the honey bee (Hymenoptera: Apidae). Annals of the Entomological Society of America 102, 881–888.

27. Wei, H., He, X.J., Liao, C.H., Wu, X.B., Jiang, W.J., Zhang, B., Zhou, L.B., Zhang, L.Z., Barron, A.B., and Zeng, Z.J. (2019). A maternal effect on queen production in honeybees. Current Biology 29, 2208–2213. e3.

28. Al-Kahtani, S.N., and Bienefeld, K. (2021). Strength surpasses relatedness–queen larva selection in honeybees. PloS ONE 16, e0255151.

29. Wagner, C.R., Mahowald, A.P., and Miller, K.G. (2002). One of the two cytoplasmic actin isoforms in Drosophila is essential. Proc Natl Acad Sci USA 99, 8037.

30. Kim, J.-G., Islam, R., Cho, J.Y., Jeong, H., Cap, K.-C., Park, Y., Hossain, A.J., and Park, J.-B. (2018). Regulation of RhoA GTPase and various transcription factors in the RhoA pathway. Journal of Cellular Physiology 233, 6381–6392.

31. Krist, M. (2011). Egg size and offspring quality: a meta-analysis in birds. Biological Reviews 86, 692–716.

32. Tetreau, G., Dhinaut, J., Gourbal, B., and Moret, Y. (2019). Trans-generational immune priming in invertebrates: current knowledge and future prospects. Frontiers in Immunology 10, 1938.

33. Plaistow, S.J., Lapsley, C.T., and Benton, T.G. (2006). Context-dependent intergenerational effects: the interaction between past and present environments and its effect on population dynamics. The American Naturalist 167, 206–215.

34. Einum, S., and Fleming, I.A. (2004). Environmental unpredictability and offspring size: conservative versus diversified bet-hedging. Evolutionary Ecology Research 6, 443–455.

35. Rollinson, N., and Hutchings, J.A. (2013). Environmental quality predicts optimal egg size in the wild. The American Naturalist 182, 76–90.

36. Rueppell, O., Kaftanouglu, O., and Page, R.E. (2009). Honey bee (Apis mellifera) workers live longer in small than in large colonies. Experimental Gerontology 44, 447–452.

37. Amdam, G.V., and Omholt, S.W. (2002). The regulatory anatomy of honeybee lifespan. Journal of Theoretical Biology 216, 209–228.

38. Rueppell, O., Aumer, D., and Moritz, R.F. (2016). Ties between ageing plasticity and reproductive physiology in honey bees (Apis mellifera) reveal a positive relation between fecundity and longevity as consequence of advanced social evolution. Current Opinion in Insect Science 16, 64–68.

39. Al-Khafaji, K., Tuljapurkar, S., Carey, J.R., and Page, R.E. (2009). Life in the colonies: Hierarchical demography of a social organism. Ecology 90, 556–566.

40. Smith, M.L., Koenig, P.A., and Peters, J.M. (2017). The cues of colony size: how honey bees sense that their colony is large enough to begin to invest in reproduction. Journal of Experimental Biology 220, 1597–1605.

41. Dixit, T., English, S., and Lukas, D. (2017). The relationship between egg size and helper number in cooperative breeders: a meta-analysis across species. PeerJ 5, e4028.

42. Summers, K., McKeon, C.S., Heying, H., Hall, J., and Patrick, W. (2007). Social and environmental influences on egg size evolution in frogs. Journal of Zoology 271, 225–232.

43. Maeno, K.O., Piou, C., and Ghaout, S. (2020). The desert locust, Schistocerca gregaria, plastically manipulates egg size by regulating both egg numbers and production rate according to population density. Journal of Insect Physiology 122, 104020.

44. Hollis, B., Koppik, M., Wensing, K.U., Ruhmann, H., Genzoni, E., Erkosar, B., Kawecki, T.J., Fricke, C., and Keller, L. (2019). Sexual conflict drives male manipulation of female postmating responses in Drosophila melanogaster. Proc Natl Acad Sci USA 116, 8437–8444.

45. Lerner, D.W., McCoy, D., Isabella, A.J., Mahowald, A.P., Gerlach, G.F., Chaudhry, T.A., and Horne-Badovinac, S. (2013). A Rab10-dependent mechanism for polarized basement membrane secretion during organ morphogenesis. Developmental Cell 24, 159–168.

46. Luo, W., Liu, S., Zhang, W., Yang, L., Huang, J., Zhou, S., Feng, Q., Palli, S.R., Wang, J., Roth, S., et al. (2021). Juvenile hormone signaling promotes ovulation and maintains egg shape by inducing expression of extracellular matrix genes. Proc Natl Acad Sci USA 118, e2104461118.

47. McDonough-Goldstein, C.E., Pitnick, S., and Dorus, S. (2021). Drosophila oocyte proteome composition covaries with female mating status. Scientific reports 11, 1–12.

48. Wheeler, D. (1996). The role of nourishment in oogenesis. Annual Review of Entomology 41, 407–431.

49. Shimada, Y., Burn, K.M., Niwa, R., and Cooley, L. (2011). Reversible response of protein localization and microtubule organization to nutrient stress during Drosophila early oogenesis. Developmental Biology 355, 250–262.

50. Wilson, M.J., Abbott, H., and Dearden, P.K. (2011). The evolution of oocyte patterning in insects: multiple cell-signaling pathways are active during honeybee oogenesis and are likely to play a role in axis patterning. Evolution & Development 13, 127–137.

51. Murphy, A.M., and Montell, D.J. (1996). Cell type-specific roles for Cdc42, Rac, and RhoL in Drosophila oogenesis. The Journal of Cell Biology 133, 617–630.

52. Hall, A. (1998). Rho GTPases and the actin cytoskeleton. Science 279, 509–514.

53. Sokolova, M., Moore, H.M., Prajapati, B., Dopie, J., Meriläinen, L., Honkanen, M., Matos, R.C., Poukkula, M., Hietakangas, V., and Vartiainen, M.K. (2018). Nuclear actin is required for transcription during Drosophila oogenesis. IScience 9, 63–70.

54. Pimm, M.L., and Henty-Ridilla, J.L. (2021). New twists in actin–microtubule interactions. Molecular Biology of the Cell 32, 211–217.

55. Denk-Lobnig, M., and Martin, A.C. (2019). Modular regulation of Rho family GTPases in development. Small GTPases 10, 122–129.

56. Amiri, E., Herman, J.J., Strand, M.K., Tarpy, D.R., and Rueppell, O. (2020). Egg transcriptome profile responds to maternal virus infection in honey bees, Apis mellifera. Infection, Genetics and Evolution 85, 104558.

57. Roth, A., Vleurinck, C., Netschitailo, O., Bauer, V., Otte, M., Kaftanoglu, O., Page, R.E., and Beye, M. (2019). A genetic switch for worker nutrition-mediated traits in honeybees. PLoS Biology 17, e3000171.

58. Baron, G.L., Raine, N.E. and Brown M.J.F. (2017). General and species-specific impacts of a neonicotinoid insecticide on the ovary development and feeding of wild bumblebee queens. Proc. R. Soc. B 284, 20170123.

59. Laidlaw, H.H., and Page, R.E. (1997). Queen Rearing and Bee Breeding (Wicwas Press).

60. Waddington, K.D. (1989). Implications of variation in worker body size for the honey bee recruitment system. Journal of Insect Behavior 2, 91–103.

61. Fang, Y., Feng, M., Han, B., Lu, X., Ramadan, H., and Li, J. (2014). In-depth proteomics characterization of embryogenesis of the honey bee worker (Apis mellifera ligustica). Molecular & Cellular Proteomics 13, 2306–2320.

62. Chen, C., Chen, H., Zhang, Y., Thomas, H.R., Frank, M.H., He, Y., and Xia, R. (2020). TBtools: an integrative toolkit developed for interactive analyses of big biological data. Molecular Plant 13, 1194–1202.

63. Xie, C., Mao, X., Huang, J., Ding, Y., Wu, J., Dong, S., Kong, L., Gao, G., Li, C.-Y., and Wei, L. (2011). KOBAS 2.0: a web server for annotation and identification of enriched pathways and diseases. Nucleic Acids Research 39, W316–W322.

64. Bindea, G., Mlecnik, B., Hackl, H., Charoentong, P., Tosolini, M., Kirilovsky, A., Fridman, W.-H., Pagès, F., Trajanoski, Z., and Galon, J. (2009). ClueGO: a Cytoscape plug-in to decipher functionally grouped gene ontology and pathway annotation networks. Bioinformatics 25, 1091–1093.

65. Szklarczyk, D., Franceschini, A., Wyder, S., Forslund, K., Heller, D., Huerta-Cepas, J., Simonovic, M., Roth, A., Santos, A., and Tsafou, K.P. (2015). STRING v10: protein–protein interaction networks, integrated over the tree of life. Nucleic Acids Research 43, D447–D452.

66. Wang, F., Flanagan, J., Su, N., Wang, L.-C., Bui, S., Nielson, A., Wu, X., Vo, H.-T., Ma, X.-J., and Luo, Y. (2012). RNAscope: a novel in situ RNA analysis platform for formalin-fixed, paraffin-embedded tissues. The Journal of Molecular Diagnostics 14, 22–29.

67. Han, B., Wei, Q., Wu, F., Hu, H., Ma, C., Meng, L., Zhang, X., Feng, M., Fang, Y., Rueppell, O., et al. (2021). Tachykinin signaling inhibits task-specific behavioral responsiveness in honeybee workers. eLife 10, e64830.

68. Vandesompele, J., De Preter, K., Pattyn, F., Poppe, B., Van Roy, N., De Paepe, A., and Speleman, F. (2002). Accurate normalization of real-time quantitative RT-PCR data by geometric averaging of multiple internal control genes. Genome Biology 3, 1–12.

69. Livak, K.J., and Schmittgen, T.D. (2001). Analysis of relative gene expression data using real-time quantitative PCR and the 2(T)(-Delta Delta C) method. Methods 25, 402–408.

